# Meta-analysis and Commentary: Preemptive Correction of Arteriovenous Access Stenosis

**DOI:** 10.1101/179580

**Authors:** Jochen G. Raimann, Levi Waldron, Elsie Koh, Gregg A. Miller, Murat H. Sor, Richard J. Gray, Peter Kotanko

## Abstract

**Background:** A recent meta-analysis by Ravani and colleagues (Ravani, P., et al., Am J Kidney Dis, 2016. 67(3): p. 446-60.) studied the effect of pre-emptive correction of arterio-venous dialysis vascular access versus deferred care, based on data from 11 trials. The authors reported a non-significant protective treatment effect of pre-emptive correction on access loss, while showing a significant protective effect on thrombosis rates conferred by pre-emptive correction. We revisit this analysis, including data extraction and effects of a heterogenous study population.

**Methods:** We repeated data extraction from all referenced publications in the meta-analysis by Ravani et al. and corrected event counts where applicable. We repeated the meta-analyses with access loss as the outcome for studies that recruited patients with arterio-venous fistulae (AVF) and grafts (AVG), respectively, using a random effects model with relative risk (RR) and risk difference (RD) of access loss as the outcomes of interest. We repeated data extraction from all referenced publications, and corrected event counts where applicable.

**Results:** Our conclusions differ from the original findings in two ways. First, after some amendment of the event counts extracted from Mayer et al. (Vascular and Endovascular Surgery 1993), we find a significant overall positive effect of pre-emptive correction on arterio-venous access loss in the overall study population [RR 0.80 (95% CI 0.64 to 0.99), RD −0.07 (95% CI −0.12 to −0.02); Figure 1]. Secondly, we highlight the impact of heterogeneous study populations on the meta-analysis. Whereas the data do not conclusively show a benefit of pre-emptive correction for arteriovenous grafts (AVG; RR = 0.87, 95% CI: 0.69 – 1.11), they show a strong protective effect for arteriovenous fistulae (AVF; RR = 0.5, 95% CI: 0.29 to 0.86).

Figure 1:
Meta-analysis of access loss, overall and by access type using risk ratio (RR) as the measure of association.

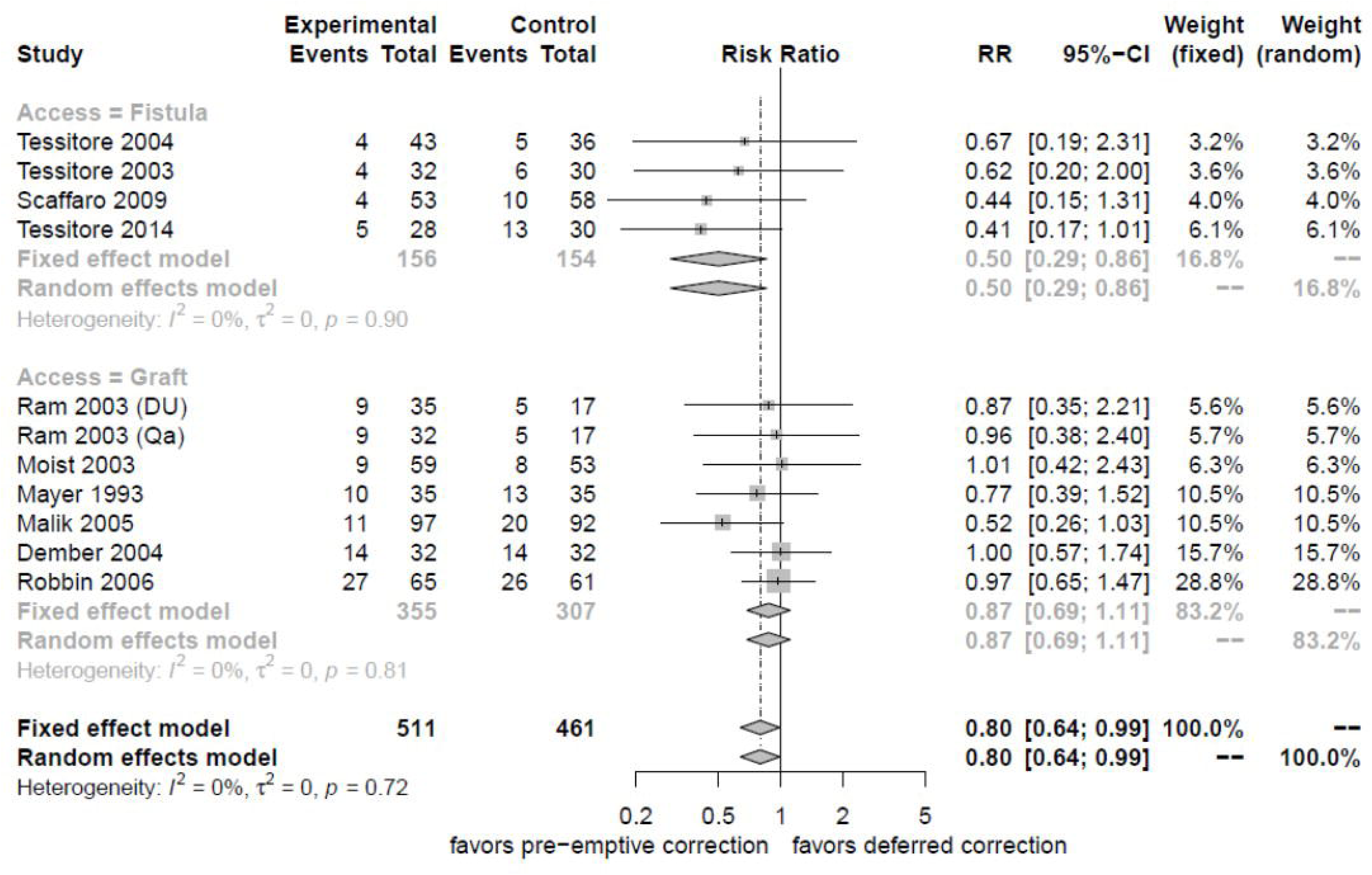

**Discussion and Conclusion:** These findings corroborate clinical arguments such as superior long-term patency of AVF and the nature of AVG failure that often involve infectious causes. The available data indicate mild or no benefit of pre-emptive correction for AVG, but strongly support tight monitoring of dialysis accesses and preemptive intervention and correction upon the slightest suspicion of access stenosis for AVF.

## Introduction

The creation and maintenance of a well-functioning vascular access is central to dialysis therapy and remains a major concern to the community. Access-related complications and failure are the most common cause of hospitalization in the dialysis population ^1^. Close monitoring by the dialysis staff and early intervention in specialized facilities has become the standard-of-care as mandated by K-DOQI, and is widely held to prolong patency and decrease the risk of adverse outcomes. Nevertheless, this approach is not universally accepted to reduce hospitalizations, access thrombosis or access loss, and thus has been the subject of an ongoing debate. Frequent access care and follow-up in free-standing offices has been shown in a large retrospective analysis to result in fewer hospitalizations, lower mortality rates, and decreased incidence of infection, resulting in lower healthcare costs to Medicare ^2^. However, the data on close observation in dialysis clinics, frequent follow-ups, and early intervention are limited to a small number of prospective studies that lack adequate power. Prospective studies of subjects receiving treatment with arterio-venous fistula (AVF), arterio-venous graft (AVG), or both, have compared rates of access loss between treatment groups receiving preemptive care, generally based on monitoring and immediate intervention, and control groups where intervention is generally performed only after clinically manifest problems develop with the access ^3-13^.

A systematic literature review and a meta-analysis recently published by Ravani et. al ^3^ strived to clarify that question. The authors investigated the effects of preemptive compared to deferred correction of AVF and AVG, respectively, on thrombosis rates, access loss, hospitalizations, procedures, infections and mortality ^3^. Preemptive care was defined as intervention following the diagnosis of stenosis using a variety of monitoring methods in the dialysis clinics. The authors systematically reviewed the literature and found 4 studies with AVF ^4-7^ and 6 studies with AVG ^8-13^. The authors reported a significant beneficial treatment effect of preemptive care on thrombosis rates. Surprisingly, the authors reported a lack of significant treatment effect on access loss, the other primary outcome [RR 0.81 (95% CI 0.65 to 1.02)]. Although Ravani and colleagues reported a 50% reduction for AVF access loss, the lack of statistical significance in the overall model has sparked a major debate in the community ^3^.

We reviewed the data extraction conducted by Ravani and colleagues, reproduced all elements of the analysis, and investigated the impact of study heterogeneity. Our findings assert that the evidence does in fact support strong benefits of preemptive correction for AVF, while being inconclusive for AVG.

## Material and Methods

We repeated all elements of the meta-analysis by Ravani *et al.* First, we repeated data extraction from all referenced publications and corrected event counts where applicable. Secondly, we repeated all meta-analyses with access loss as the outcome. Finally, we used Monte Carlo simulation to demonstrate the effect on meta-analysis synthesis of combining studies of AVG and AVF patients in varying proportions. Analyses were done in R version 3.2.2 (R Foundation for Statistical Computing; Vienna, Austria) ^14^ using the packages *meta, metafor* and *lme*. A p-value less than 5% was considered significant.

### Meta-Analysis

We reviewed all publications used in the meta-analysis of Ravani and colleagues ^3^, extracting event counts using the methods reported in the above Cochrane review ^15^. This meta-analysis focuses on the outcome of access loss. All studies consider the comparison a treatment group receiving preemptive access correction to a control group receiving deferred correction. Each individual study comprises either a) AVF or b) AVG patients, and no study combines these two patient groups. Random and fixed-effects meta-analysis were performed using risk ratio (RR) and risk difference (RD) as effect measures. In addition to combining both access types in the same fashion as Ravani and colleagues ^3^, we analyzed the two dialysis access types separately. Statistical heterogeneity was quantified by Cochran’s Q and the magnitude of variation, I^2^. The two access types were tested for subgroup differences using meta-regression using the DerSimonian-Laird and the Hartung-Knapp method.

### Bootstrap / Monte Carlo Simulation

Total sample size and numbers of events from experimental and control arms of the 11 studies were extracted from Figure 3 of Ravani *et al.* ^3^. This meta-analysis included four studies of fistula access type and seven studies of graft access type. The event probabilities in each arm of each study (π_i,control_, π_i,experimental_) were calculated as the ratio of the number of events to the total sample size of that arm. The following steps were performed for the simulation, for numbers of fistula studies NS_fistula_ between zero and 11:

1. **Bootstrap step:** randomly select NS_fistula_ studies (with replacement) from the four fistula studies, and (11 -NS_fistula_) studies (with replacement) from the 7 graft studies, for a total of 11 studies in the bootstrap sample
2. **Monte Carlo step:** for each study *i*, simulate the number of events x_i,control_ in the control arm from a random binomial distribution with event probability π_i,control_, and n_i,control_ trials. Do the same for each experimental arm, simulating the number of events x_i,experimental_ using event probability π_i,experimental_, and n_i,experimental_ trials.
3. **Meta-analysis step:** Using these simulated numbers of events (x_i,control_, x_i,experimental_) and sample sizes (n_i,control_, n_i,experimental_), perform random-effects meta-analysis of RR or OR equivalently to Figure 3 of Ravani *et al.*
4. Repeat steps 1-3, 100 times
5. Calculate the mean OR/RR and upper/lower confidence intervals from step 3 These steps were repeated with NS_fistula_ = 0, 1, 2,…,11 to simulate the effect of mixing fistula and graft access type studies in varying proportions.

## Results

### Replication of Ravani et al. Meta-analysis

We repeated data extraction of all reports included in the meta-analysis of Ravani *et al.* and noted a data transfer error in the number of events for the study by Mayer et al. ^10^. Ravani et al. stated an equal number of events (n=10) in control (deferred correction) and experimental group (pre-emptive correction) ^3^, while the actual numbers would correctly be reported as 10 for the experimental and 13 for the control group ^10^. After correction of this event count error, the pooled meta-analyses of AVF and AVG patients showed a significant protective effect of preemptive correction [RR 0.80 (95% CI 0.64 to 0.99), RD −0.07 (95% CI −0.12 to −0.02)].

### Subgroup Analysis

The meta-analysis presented with non-significant heterogeneity when quantified with either I^2^ (I^2^=0%) or as residual heterogeneity (QE=3.6; P=0.94). The meta-regression showed a non-significant difference when the DerSimonian-Laird method was employed (QM=3.44; P=0.06), while it was significant when the Hartung-Knapp method was used (F=8.6; P=0.02).

However, most importantly, the protective effect of preemptive correction differed by access type. The intervention approximately halves the risk of access loss for patients with AVF [RR 0.50 (95% CI 0.29 to 0.86) and RD −0.10 (95% CI −0.17 to −0.02)], but has a weaker and non-significant protective effect among the studies of patients with AVG [RR 0.80 (95% CI 0.64 to 0.99) and RD −0.07 (95% CI −0.12 to −0.02)]; **Figure 1** and **Figure 2**. Of note, based on the Baujat plot the effect towards the Null in the overall model seems to a large extent be driven by the data from the study of Robbin and colleagues (**Figure 3**) ^13^.

**Figure 2:**
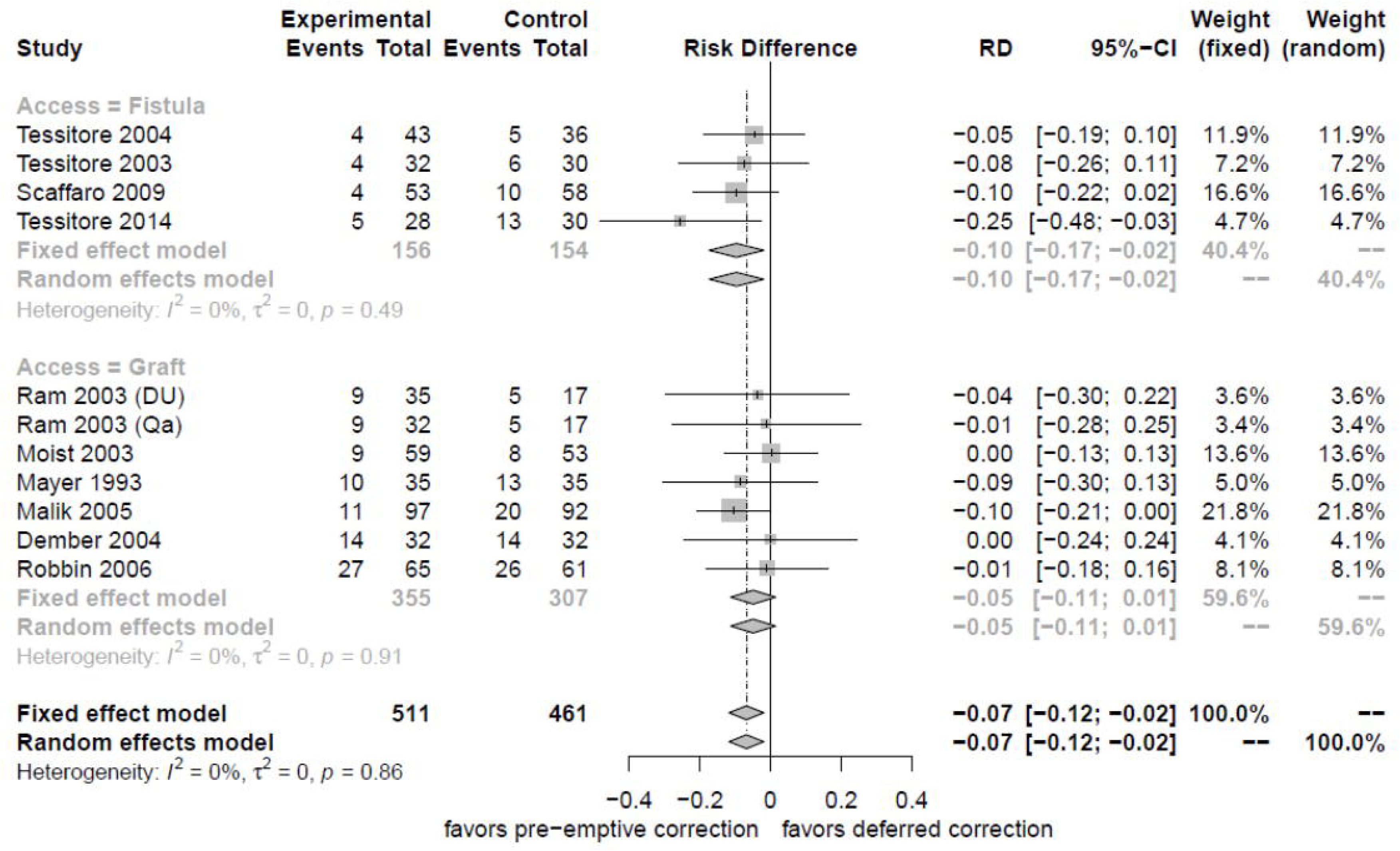
Meta-analysis of access loss, overall and by access type using risk difference (RD) as the measure of association.

**Figure 3:**
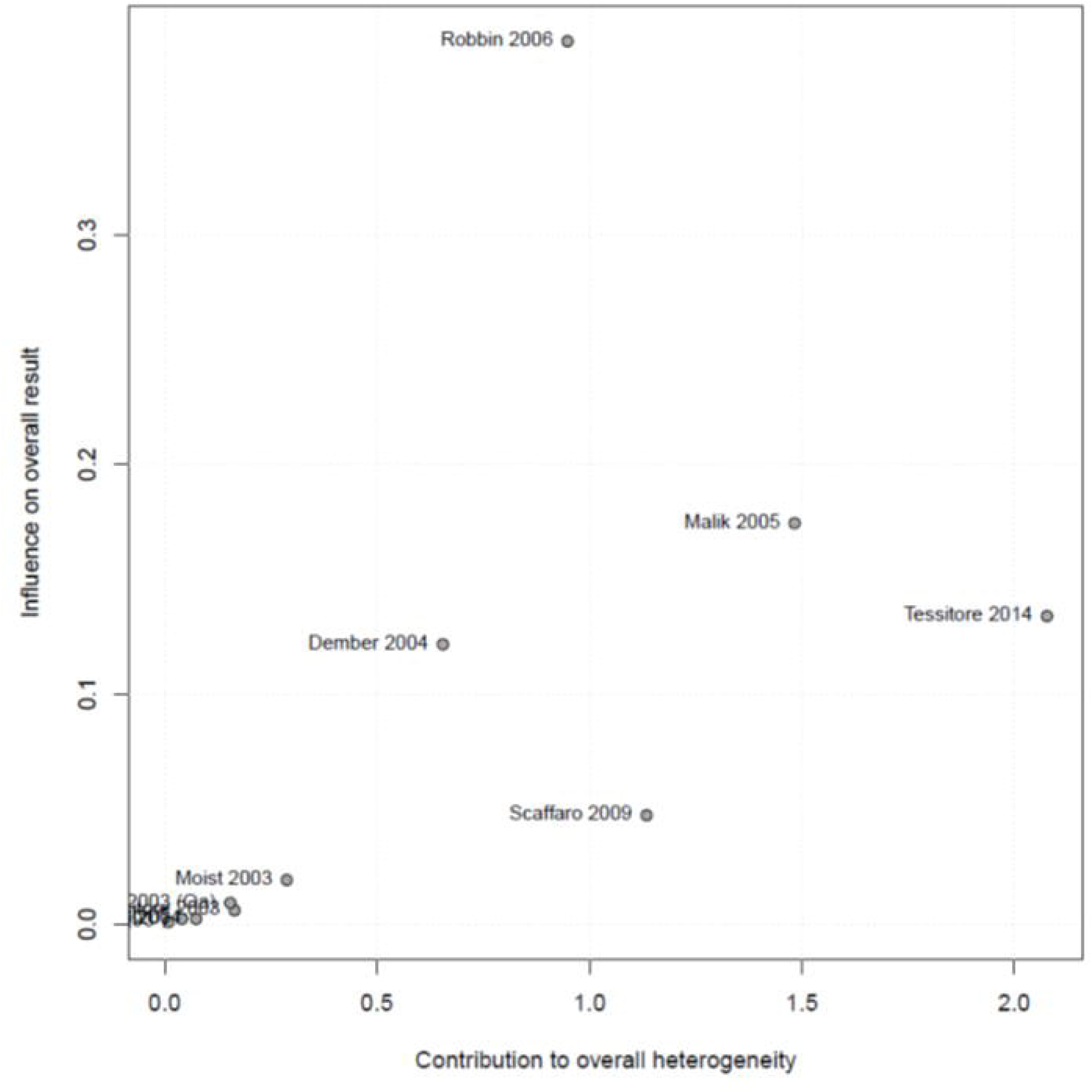
Baujat plot of the influence of each included study on the overall result as a function of the respective contribution to the overall heterogeneity.

### Simulating the Effect of Heterogeneous Populations

We employed a hybrid bootstrap / Monte Carlo simulation to demonstrate the impact of mixing heterogeneous populations in the meta-analysis. When mixing heterogeneous patient populations, such as patients with graft access type stenosis where pre-emptive correction has mild or no effect, with patients with fistula access type stenosis where pre-emptive correction has a strong protective effect, the results depend on the arbitrary proportions of the two populations included. This proportion is arbitrary because it depends on the number of studies of each population that have been published. We simulated meta-analyses based on the sample sizes and event probabilities of the original studies included in the meta-analysis by Ravani *et al.* ^3^, but where we manipulated the proportion of studies of fistula and graft access type stenosis. In other words, the number of studies was held constant at 11, but the number based on fistula access-type studies was varied from 0 to 11. Due to the greater effectiveness of preemptive correction on fistula access type stenosis, meta-analysis of intervention effectiveness produces results that are not statistically significant when zero to three fistula-type studies are included, but reach statistical significance for four or more fistula-type studies. The synthesized risk ratio is 0.86(sub exact number) when all studies are graft access type, and 0.50(sub exact number) when all studies are fistula access type (**Figure 4**). This exercise demonstrates the arbitrariness of meta-analysis that synthesizes heterogeneous populations where treatment effectiveness differs.

**Figure 4:**
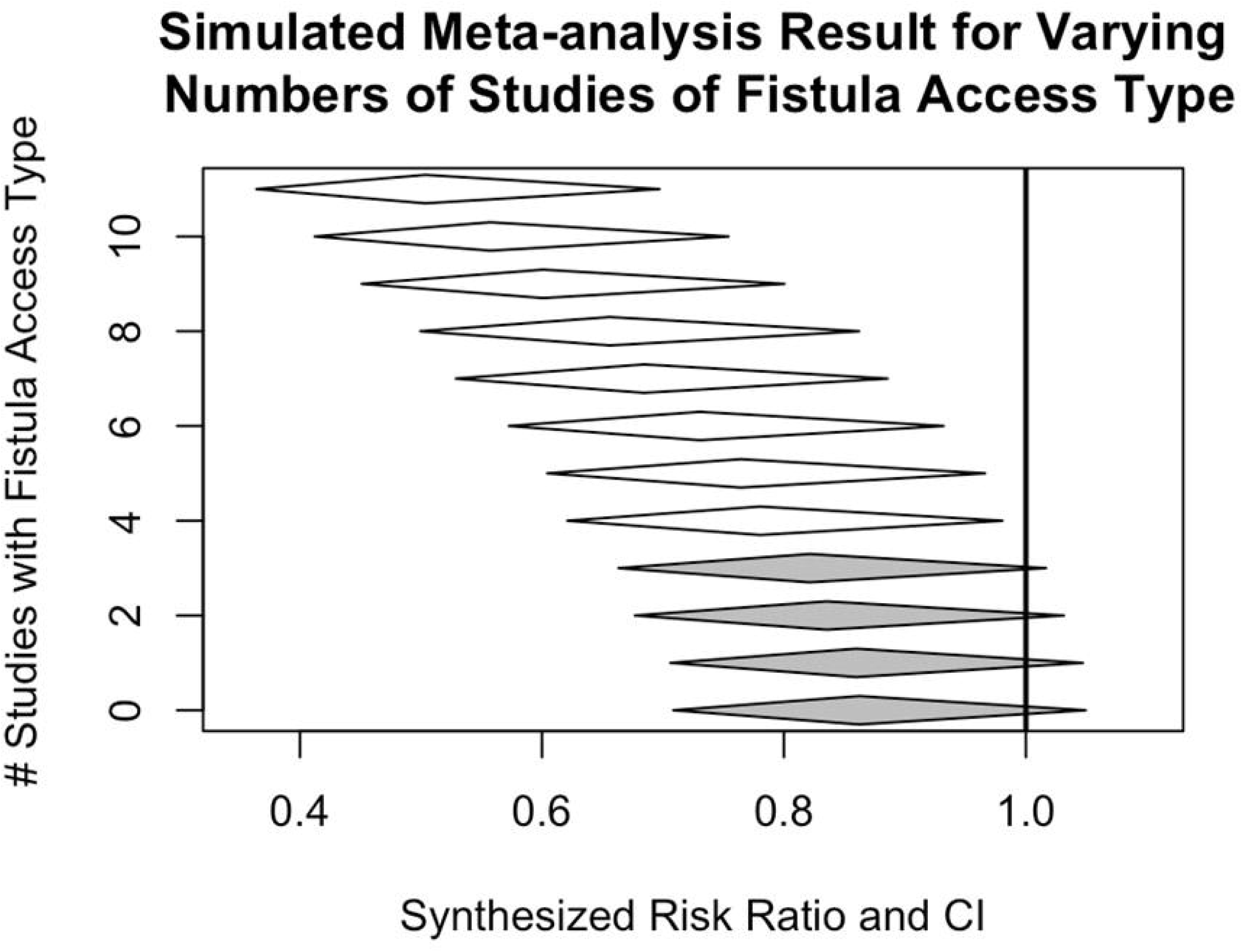
Bootstrap / Monte Carlo simulation of combining graft and fistula access type studies in varying proportions. In each iteration of the simulation, 11 studies drawn (with replacement) from the original 11 studies of access loss considered by Ravani *et al.*, drawing from the fistula studies the number of simulated studies specified on the y-axis, and the remainder of the 11 from the graft access type studies. Numbers of events were simulated using the same event probabilities as observed in the original studies. Diamonds represent the average risk ratio and confidence intervals synthesized by random effects meta-analysis and averaged across 100 simulations. Those failing to reach significance at alpha = 0.05 are highlighted in grey (0 to 3 fistula studies). Four fistula studies, the number included in the meta-analysis by Ravani *et al.*, happens to be the number at which the meta-analysis crosses the threshold of significance. This simulation highlights the arbitrariness of the meta-analysis result when synthesizing studies of both access types in arbitrary proportions.

## Discussion

### Statement of Principal Findings

We revisited the meta-analysis conducted by Ravani and colleagues with the Cochrane Database Collaboration and found, after correction of a data transfer error by the authors, a statistically significant protective effect of preemptive correction on access survival in patients receiving dialysis with either AVG or AVF [RR 0.80 (95% CI 0.64 to 0.99) and RD −0.07 (95% CI −0.12 to −0.02); **Figure 1** and **Figure 2**]. More importantly, we want to highlight that preemptive correction is much more effective for patients with AVF [RR 0.50 (95% CI 0.29 to 0.86) and RD −0.10 (95% CI −0.17 to −0.02)]. We demonstrate by simulation the arbitrariness of results based on a synthesis of AVG and AVF access type studies, noting that none of the individual studies included in this meta-analysis themselves mixed AVG and AVF access types. Both populations are conceptually different with different dynamics in terms of access patency and long-term survival. Thus pooling these two populations for such an investigation of access-related research questions needs to be avoided. **Figure 1** and **Figure 2** support the effectiveness of preemptive correction on access survival and the consequential benefit for the individual patient. Given the differences between AVG and AVF subset analyses, however, this protective effect is undoubtedly more pronounced for AVF (RR 0.41 for AVF versus 0.97 for AVG).

### Discussion of Results in the light of other studies

Meta-analysis studies must be considered with appropriate caution because they can have significantly different conclusions than subsequent large randomized controlled trials of the same topic ^16^. Study heterogeneity can harm the validity of meta-analysis and result in disagreement with gold-standard large randomized controlled trials ^16^. This is because there may be biological reasons, statistical evidence, or both, showing that “the studies included in the meta-analysis have in fact measured somewhat different things, so that a combined estimate cannot be meaningful unless additional, doubtful assumptions are made.” ^17^, There is sound reason to expect pre-emptive correction to be more important for fistula than for grafts access type, given the higher incidence of early onset failure in concert with the substantially superior long-term survival of AVF compared to AVG. In this context the statement by Bailar ^17^ appears perfectly accurate and it may indeed be hypothesized that high-frequent follow-ups by the vascular surgeon or interventionist shortly after access creation contributes to its survival during the initial high-risk period. AVG accesses tend to fail at a later time point in time, often with severe comorbidities and infections involved, that result in severe complications with access resulting in substantially shorter long-term survival rates for AVG, thus an explanation for the lack of a significant treatment effect of preemptive correction could be the overall condition of the access (and possibly also the patient). For this reason, and the consistent observed difference in treatment effectiveness, we do not believe it is justifiable to present a single estimate of effectiveness that synthesizes AVG and AVF access type. This is consistent with the 11 individual studies analyzed, none of which reported estimates that combined the two access types.

### Clinical Implication

Stenoses of accesses are the most important predictors of access failure. Early detection of stenoses and adequate counteraction that restores the fistula blood flow will increase the probability of reducing the risk of thrombosis, stenosis progression and ultimately access failure and loss. The importance of constant unobstructed flow has been known since the definition of Virchow’s triad (stasis, vessel wall injury and hypercoagulability). Given that stasis is a main component of thrombosis, the benefits of pre-emptive treatment for stenosis in the context of dialysis access are supported by the data of the meta-analysis for AVF studies showing a 50% risk reduction. Given the different nature of complications in AVGs (often involving inflammatory processes) mainly occurring at a later point in time, the less pronounced effect of preemptive correction for AVGs appears also reasonable.

The conclusions made by Ravani et al and subsequently by interpretations published online, diminish the significance of state-of-the art access monitoring techniques and the need to act at the slightest suspicion of compromise to the intra-access blood flow. In addition, if the subsequent argument is made that that intervention should then be reserved for when the accesses thrombose, studies have shown, such as the one by Urbanes and colleagues ^18^,a 5-fold higher risk of severe adverse events for thrombectomies compared to fistulograms with angioplasties. Last, the potential to reduce costs to the healthcare system ^2^ may be lost if the reader or members of the medical community become misguided by conclusions drawn by Ravani et al. or its subsequent published interpretations.

## Conclusion

Whereas Ravani *et al.* argued that the small sample size of AVF studies renders the outcome of the overall meta-analysis insufficiently conclusive, we show that the four relatively small AVF studies are on their own sufficient for statistically significant evidence of a strong protective effect of preemptive correction of access stenosis. The larger number and sample size of AVG studies, conversely, provide inconclusive evidence of a weaker protective effect. We believe this meta-analysis corroborates tight monitoring of dialysis accesses and preemptive intervention and correction upon the slightest suspicion of access stenosis for AVF. Although the evidence for a protective effect of AVG is inconclusive, we would recommend frequent follow-ups and continued preemptive correction, particularly in the initial time period following access creation, until large prospective, randomized and adequately powered study data are available for both types of accesses.

## References

1. Saran R, Li Y, Robinson B, Ayanian J, Balkrishnan R, Bragg-Gresham J, et al. US Renal Data System 2014 Annual Data Report: Epidemiology of Kidney Disease in the United States. American journal of kidney diseases : the official journal of the National Kidney Foundation. 2015;65(6 Suppl 1): A7.

2. Dobson A, El-Gamil AM, Shimer MT, DaVanzo JE, Urbanes AQ, Beathard GA, Litchfield TF. Clinical and economic value of performing dialysis vascular access procedures in a freestanding office-based center as compared with the hospital outpatient department among Medicare ESRD beneficiaries. Seminars in dialysis. 2013;26(5): 624–632.

3. Ravani P, Quinn RR, Oliver MJ, Karsanji DJ, James MT, MacRae JM, et al. Preemptive Correction of Arteriovenous Access Stenosis: A Systematic Review and Meta-analysis of Randomized Controlled Trials. American journal of kidney diseases : the official journal of the National Kidney Foundation. 2016;67(3): 446–460.

4. Scaffaro LA, Bettio JA, Cavazzola SA, Campos BT, Burmeister JE, Pereira RM, et al. Maintenance of Hemodialysis Arteriovenous Fistulas by an Interventional Strategy. Journal of Ultrasound in Medicine. 2009;28(9): 1159–1165.

5. Tessitore N, Bedogna V, Poli A, Lipari G, Pertile P, Baggio E, et al. Should current criteria for detecting and repairing arteriovenous fistula stenosis be reconsidered? Interim analysis of a randomized controlled trial. Nephrology, dialysis, transplantation : official publication of the European Dialysis and Transplant Association -European Renal Association. 2014;29(1): 179–187.

6. Tessitore N, Lipari G, Poli A, Bedogna V, Baggio E, Loschiavo C, et al. Can blood flow surveillance and pre-emptive repair of subclinical stenosis prolong the useful life of arteriovenous fistulae? A randomized controlled study. Nephrology, dialysis, transplantation : official publication of the European Dialysis and Transplant Association -European Renal Association. 2004;19(9): 2325–2333.

7. Tessitore N, Mansueto G, Bedogna V, Lipari G, Poli A, Gammaro L, et al. A prospective controlled trial on effect of percutaneous transluminal angioplasty on functioning arteriovenous fistulae survival. Journal of the American Society of Nephrology : JASN. 2003;14(6): 1623–1627.

8. Dember LM, Holmberg EF, Kaufman JS. Randomized controlled trial of prophylactic repair of hemodialysis arteriovenous graft stenosis. Kidney international. 2004;66(1): 390–398.

9. Malik J, Slavikova M, Svobodova J, Tuka V. Regular ultrasonographic screening significantly prolongs patency of PTFE grafts. Kidney international. 2005;67(4): 1554–1558.

10. Mayer DA, Zingale RG, Tsapogas MJ. Duplex Scanning of Expanded Polytetrafluoroethylene Dialysis Shunts: Impact on Patient Management and Graft Survival. Vascular Surgery. 1993;27(9): 647–658.

11. Moist LM, Churchill DN, House AA, Millward SF, Elliott JE, Kribs SW, et al. Regular monitoring of access flow compared with monitoring of venous pressure fails to improve graft survival. Journal of the American Society of Nephrology : JASN. 2003;14(10): 2645–2653.

12. Ram SJ, Work J, Caldito GC, Eason JM, Pervez A, Paulson WD. A randomized controlled trial of blood flow and stenosis surveillance of hemodialysis grafts. Kidney international. 2003;64(1): 272–280.

13. Robbin ML, Oser RF, Lee JY, Heudebert GR, Mennemeyer ST, Allon M. Randomized comparison of ultrasound surveillance and clinical monitoring on arteriovenous graft outcomes. Kidney international. 2006;69(4): 730–735.

14. R Development Core Team. R: A language and environment for statistical computing. Vienna, Austria: R Foundation for Statistical Computing, 2015.

15. Ravani P, Quinn RR, Oliver MJ, Karsanji DJ, James MT, MacRae JM, et al. Pre-emptive correction for haemodialysis arteriovenous access stenosis. The Cochrane database of systematic reviews. 2016(1): CD010709.

16. LeLorier J, Gregoire G, Benhaddad A, Lapierre J, Derderian F. Discrepancies between meta-analyses and subsequent large randomized, controlled trials. The New England journal of medicine. 1997;337(8): 536–542.

17. Bailar JC, 3rd. The promise and problems of meta-analysis. The New England journal of medicine. 1997;337(8): 559–561.

18. Urbanes AQ. Dialysis access procedures in the outpatient setting: risky? Journal of vascular and interventional radiology : JVIR. 2013;24(12): 1787–1789.

